# Human inborn errors of long-chain fatty acid oxidation show impaired inflammatory responses to TLR4-ligand LPS

**DOI:** 10.1101/2023.08.30.555512

**Authors:** Signe Mosegaard, Krishna S. Twayana, Simone W. Denis, Jeffrey Kroon, Bauke V. Schomakers, Michel van Weeghel, Riekelt H. Houtkooper, Rikke K. J. Olsen, Christian K. Holm

**Affiliations:** Research Unit for Molecular Medicine, Department of Clinical Medicine, Aarhus University and Aarhus University Hospital, Aarhus, Denmark; Department of Biomedicine, Aarhus Research Center for Innate Immunology, Aarhus University, Aarhus, Denmark; Laboratory Genetic Metabolic Diseases, Amsterdam UMC, University of Amsterdam, Amsterdam, The Netherlands; Amsterdam Gastroenterology, Endocrinology, and Metabolism, Amsterdam, The Netherlands; Amsterdam Cardiovascular Sciences, Amsterdam, The Netherlands; Department of Experimental Vascular Medicine, Amsterdam UMC, University of Amsterdam, Amsterdam, The Netherlands; Core Facility Metabolomics, Amsterdam University Medical Centers, University of Amsterdam, Amsterdam, The Netherlands; Emma Center for Personalized Medicine, Amsterdam UMC, Amsterdam, The Netherlands

**Keywords:** Long-chain fatty acid oxidation disorders, immunometabolism, lipopolyssacharide (LPS), toll-like receptor 4 (TLR4)

## Abstract

Stimulation of mammalian cells with inflammatory inducers such as lipopolysaccharide (LPS) leads to alterations in the activity of central cellular metabolic pathways. Interestingly, these metabolic changes seem to be important for the subsequent release of pro-inflammatory cytokines. This has become particularly clear for enzymes of the tricarboxylic acid (TCA) cycle such as succinate dehydrogenase (*SDH*). LPS leads to inhibition of SDH activity and accumulation of succinate to enhance the LPS-induced formation of IL-1β. If enzymes involved in beta-oxidation of fatty acids are important for sufficient responses to LPS is currently not clear.

Using cells from various patients with inborn fatty acid oxidation disorders, we report that disease-causing deleterious variants of Electron Transfer Flavoprotein Dehydrogenase (*ETFDH*) and of Very Long Chain Acyl-CoA Dehydrogenase (*ACADVL*), both cause insufficient responses to stimulation with LPS. The insufficiencies included reduced TLR4 expression levels, impaired TLR4 signaling, and reduced or absent induction of pro-inflammatory cytokines such as IL-6. The insufficient responses to LPS were reproduced in cells from normal healthy controls by targeted loss-of-function of either *ETFDH* or *ACADVL,* supporting that the deleterious *ETFDH* and *ACADVL* variants cause the attenuated responses to LPS.

*ETFDH* and *ACADVL* encode two distinct enzymes both involved in fatty acid beta-oxidation, and patients with these deficiencies cannot sufficiently metabolize long-chain fatty acids. With this report, we therefore provide genetic evidence from two genetically distinct but phenotypically similar human metabolic diseases, that genes important for beta-oxidation of long-chain fatty acids are also important for inflammatory responses to LPS.

## INTRODUCTION

Metabolic pathways are crucial for the proper functioning of the cell and are involved in various processes such as energy production, biosynthesis and degradation of molecules^1^. It has now become clear that these metabolic pathways also play critical roles in the regulation of immune responses^2–4^. For example, pharmacological inhibition of glycolysis also inhibits lipopolysaccharide (LPS)-induced formation of the pro-inflammatory cytokine IL-1^5^. Intriguinly, the dependency on glycolysis for the response to LPS is based on stabilization of Hypoxia-inducible Factor 1-alpha (HiF1α) by an LPS-induced increment in succinate levels^5^. Later, the increment in succinate was demonstrated to be dependent on another metabolite itaconate derived from aconitate^6^. These discoveries were among the first to demonstrate that enzymes and metabolites of the tricarboxylic acid (TCA) cycle impacted inflammatory responses to LPS. Fatty acid oxidation (FAO) also intersects with inflammatory responses and has been associated with development of anti-inflammatory phenotypes of macrophages. Indeed, induction of classical differentiation of macrophages through stimulation with LPS and Interferon gamma (IFNγ) induce an increase in glycolytic flux. By contrast, alternative activation induced by IL-4 seemed to promote an increased reliance on FAO^7,8^. In line,, forced increases in FAO in adipocytes seems to limit the pro-inflammatory phenotype induced by treatment with palmitate^9^. This link as recently been strengthened as stimulation of murine macrophages with LPS is now demonstrated to lead to an increase in FAO that involves stabilization of the mitochondrial fatty acid transporter carnitine palmitoyl transferease 1a (CPT1a)^10^. If FAO or central FAO enzymes are important for the pro-inflammatory release of cytokines in the response to stimuli like LPS is not clear. Furthermore, observations of such connection in human disease have not yet been made.

Disorders of long-chain fatty acid oxidation (lcFAOD) were first described in the 1970’s, and since, inherited defects have been described for most of the enzymes, transporters, and other facilitating proteins involved in FAO^11,12^. lcFAODs are recessively inherited and lead to impaired energy production and various clinical including cardiac- and skeletal muscle myopathies. The most common lcFAOD is VLCAD deficiency (VLCADD), which is caused by genetic variants in *ACADVL,* and has a worldwide prevalence of 1:31.500-1:94.569^13^. VLCADD can present with mild or severe disease, or even remain asymptomatic for years. A less frequent lcFAOD is multiple acyl-CoA dehydrogenation deficiency (MADD or glutaric aciduria type II (GAII)). Similar to patients with VLCADD, patients with MADD vary in their clinical presentation. Mildly affected patients present similarly to VLCADD with mainly muscle-related phenotypes such as exercise intolerance, weakness and pain. Severe and neonatal onset MADD is multisystemic and can be lethal. Most patients with MADD harbor genetic variants in genes encoding the flavoproteins ETF or ETF-QO^14^.

It is known from clinical practice that infections are a disease trigger in lcFAODs^15–17^, and that inflammation and infections can cause metabolic decompensation, rhabdomyolysis and even trigger multi-organ failure and death^14,18–20^. In humans, only a few studies describe immune responses in lcFAODs. Those studies show an inflammatory dysregulation both during symptomatic and non-symptomatic periods^17,21,22^.

Using samples from two MADD and four VLCADD patients, we here report that patient cells with the underlying deleterious variants in *ETFDH* and *ACADVL* display highly impaired responses to stimulation with LPS. This included reduced expression of TLR4, impaired TLR4 signaling, and reduced or absent induction of cytokines such as IL-6. Our study thereby delivers genetic evidence using patient-derived cells, that enzymes important for beta-oxidation of long-chain fatty acids are also critical for effective responses to stimulation with LPS.

## MATERIALS AND METHODS

### Patient and control material

Human primary fibroblasts were derived from patients previously diagnosed with severe or mild VLCADD or severe MADD, based on clinical, biochemical, and genetic data (Supplementary Materials Table 1). Control dermal fibroblasts were obtained from healthy individuals. All the cells were de-identified, and the inclusion of these samples in this study is according to the Dutch and Danish ethical committee regulations.

### Cell culture and reagents

Dermal fibroblasts from VLCADD patients (severe; P5, P6, P7, P8, and mild; P3 and P4) and controls (C3, C4, C5 and C6) were cultured until confluent with Ham’s F-10 nutrient mixture (Gibco) supplemented with L-glutamine (Bio-Whittaker), 10% fetal bovine serum (FBS) (Bio-Whittaker), 100 U/ml penicillin, 100μg/ml streptomycin (Life Technologies), and 250 ng/ml Fungizone (Life Technologies) in a humidified atmosphere with 5% CO_2_ at 37 °C. Dermal fibroblasts from MADD (P1 and P2) and controls (C1 and C2) were cultured in DMEM (Merk, D6429) supplemented with 10% heat inactivated fetal bovine serum (Merk, F9665), 200 IU/mL−1 penicillin, 100 μg/mL−1 streptomycin and 600 μg/mL−1 glutamine, hereafter termed complete DMEM. When confluent cells were seeded in triplicates in xx well plates (xx,000 cells/well) with equal confluency for 24 hours. After 24 hours, medium was replaced with Ham’s F-10 nutrient mixture, 1% FBS, as described above, with 0 ng/mL or 400 ng/mL lipopolysaccharide (LPS) (Serotype EH100 (RA) (TLRGRADE®) (Ready-to-Use), Alexis Biochemicals) or for MADD (Invivogen (tlrl-b5lps)), or 2 µg/mL poly(I:C) (Invivogen (trl-pic-5). After 24 hours incubation, medium was collected and frozen at -80°C. Cells were harvested using trypsin, and washed twice with PBS, before cell pellets were frozen at -80°C. LPS concentration and duration of incubation were chosen based on published studies and several test experiments^23–25^. Cells were regularly tested for mycoplasma contamination by sequencing from GATC Biotech (Germany). *ETFDH* siRNA (h):sc-89048 was from Santacruz biotechnology. For knockout, *ETFDH* guide RNA (ETFDH_g1, SYNTHEGO) used had sequence of GACCAUCUUGUAGCACAUAG. DuoSet ELISA Development systems from R & D were used for measurement of human IL-6 (DY206), human IL 1β (DY201), CXCL10 (DY266), and IL-10 (DY217B).

### Long-chain fatty acid oxidation flux assay

VLCADD (severe; P5, P6, P7, P8, and mild; P3 and P4) and control (C4, C5 and C6) dermal fibroblasts were seeded in 48-well plates, and the oxidation fluxes were performed in duplicates and measured at 37°C by the production of ^3^H_2_O from [9,10– ^3^H(N)]-oleic acid as previously described^26^. FAO flux in MADD and control cells was determined by measuring radiolabelled H_2_O from [9,10-^3^H(N)]-palmitic acid. FAO flux was assayed for controls, *ETFDH* mutant cells, and *ETFDH* silenced control fibroblasts. 100 µM Etomoxir (ETO), a small molecule inhibitor of CPTI, was used as positive control for FAO reduction. Cells seeded in 24 well plates (50,000 cells/well) were either transfected with siETFDH for 72 h, or pretreated with 100 µM ETO for 12 h, or directly proceeded for FAO measurement in case of controls and *ETFDH* mutant cells. Briefly, cells were incubated for 6 h with complete DMEM containing radiolabelled [9,10-^3^H] palmitic acid. Supernatant was then used to determine radioactivity by liquid scintillation counting. Measurement was done at 37°C and FAO flux was expressed in nmol palmitate/h/mg protein.

### Enzyme-linked immunoassay (ELISA)

Cytokines in the patient and control dermal fibroblast culture supernatants were measured using enzyme-linked immunosorbent assay (ELISA) for quantitative detection of IL-1β (R&D Systems Cat. No. DY201), tumour necrosis factor TNF-α (BioLegend® Cat. No. 430204), IL-6 (BioLegend® Cat. No. 430504 and R&D Systems Cat. No. DY206), IL-8 (Thermo Scientific 88-8086), IL-10 (BioLegend® Cat. No. 430604 and R&D Systems Cat. No. DY217B), and CXCL10 (R&D Systems Cat. No. DY266), all according to the manufacturer’s instructions. The plates were coated with capture antibody and after overnight incubation at 4 °C, the wells were washed (PBS + 0.05% Tween) and the samples and standard were incubated for 2 hours (shaking at room temperature). Next, the samples were aspirated, wells were washed, antibody was added and detected using Avidin-HRP. To visualize, tetramethylbenzidine (TMB) was added for 5-15 min and the enzyme substrate reaction was terminated by adding acetic acid and measured at 450 nm.

### RNA analysis and QPCR analysis

In MADD and control fibroblasts, RNA from dermal fibroblasts cell pellets were extracted using High Pure RNA Isolation Kit (Roche) using manufacturer’s instructions. RNA quantification and quality assessment was done using Nanodrop spectrometry (Thermo Fisher Scientific). Gene expression was determined by real-time quantitative PCR (Taqman Gene Expression Assay, Gene Assay IDs in Supplementary Materials Table 2), using TaqMan detection systems (Applied Biosciences). mRNA levels were determined using Taqman RNA-to-Ct 1-step Kit (Applied Biosystems) following manufacturer’s guidelines.

RNA from VLCADD and control fibroblast cell pellets were isolated by using TRI Reagent® (Sigma Aldrich) according to the manufacturer’s instructions. RNA was quantified using the NanoDrop 2000 spectrophotometer (Thermo Fisher Scientific). 1 μg of RNA was pre-treated with gDNA Wipeout Buffer and reverse-transcribed to cDNA according to the manufacturer’s instructions using QuantiTect Reverse Transcription Kit (QIAGEN). Quantitative gene expression analysis was performed using the LightCycler®480 SYBR Green I Master (Roche) and measured using the LightCycler®480 Instrument II (Roche). Relative expression was calculated by LinRegPCR^27^ and the N0 values were normalized to the geometric mean of reference genes beta-actin, GAPDH and 36B4. Primers directed toward the specific genes and sequences are listed in the Supplementary Material, Table 3.

### Western blot

Cells lysate was collected in 100 μL of ice-cold Pierce RIPA lysis buffer (Thermo Scientific) supplemented with 10 mM NaF, 1x complete protease cocktail inhibitor (Roche) and 5 IU mL−1 benzonaze (Sigma), respectively. Protein concentration was determined using a BCA protein assay kit (Thermo Scientific). Protein denaturation before separation was done by treating the cell lysates for 3 min at 95°C in presence of 1x XT Sample Buffer (BioRad) and 1x XT reducing agent (BioRad). 15 to 30 µg of protein sample was processed for separation through SDS PAGE on 4–20% Criterion TGX precast gradient gels (BioRad). Gel run consisted of 15 min run at 70V followed by 45 min run at 100V. A dry protein transfer onto PVDF membrane (BioRad) was then performed using a Trans-Blot Turbo Transfer system for 7 min. In order to prevent nonspecific binding, membranes were blocked for 1 h in 5% skim-milk (Sigma Aldrich) at room temperature in PBS supplemented with 0.05% Tween-20 (PBST). Blocked membranes were cut according to desired protein molecular weight and incubated overnight at 4°C in specific primary antibodies prepared in PBST (diluted 1:1000 unless mentioned). The one used for this study were: anti-TLR4 (sc-293072, Santa Cruz Biotechnology), anti-IRAK1 (#4504, CST), anti-P-IRAK1(# PA5-38633, Thermo Fischer Scientific), anti-IκBα (#4814, CST), anti-Phospho-IκBα (#2859, CST), anti-NF-κB p65 (#8242, CST), anti-Phospho NF-κB p65(#3033, CST), anti-ETFDH (sc-515202, Santa Cruz Biotechnology), anti-JunB (#3753, CST), anti-cJun (#9165, CST), anti-FRA1(#5281, CST), anti Phospho FRA1 (#3880, CST), and anti-Vinculin (#13901, CST). Membranes were then washed three times in PBST and incubated in secondary antibodies; peroxidase-conjugated F(ab)2 donkey anti-rabbit IgG (H + L) (1:10,000) (Jackson ImmunoResearch) prepared in 1% PBST skim milk. It was then followed by three times membranes washing in PBST and exposure to either the SuperSignal West Pico PLUS chemiluminescent substrate or the SuperSignal West Femto maximum sensitivity substrate (ThermoScientific) and Images were captured using an ImageQuant LAS4000 mini-Imager (GE Healthcare).

### Short-interfering RNA (siRNA)-mediated knockdown

For short interfering RNA experiments, cells seeded in 12 well plate (125,000 cells per well) were transfected with 80 pmol of gene specific siRNA (human *ETFDH* (sc-89048), human *JunB* (sc-35726), human *MTOR* (sc-35409) or control siRNA (sc-37007), diluted in antibiotic free DMEM, using Lipofectamine RNAi Max as per manufacturer’s instructions for 72 h. LPS 400 ng/ml stimulation was given after siRNA incubation without removing media.

### CRISPR/Cas9 knockout of *ETFDH* and activation of *JunB*

Control dermal fibroblasts (300,000 to 800,000 cells), resuspended in 20 µl Opti-MEM were electroporated with 1.5 nmol *ETFDH* guide RNA and 1.2 µl Cas9 enzyme. Cells were then seeded in prewarmed DMEM and grown until confluency for 2 to 3 days. These once knocked out cells were again collected and electroporated with similar composition of guide RNA and Cas9. Double knocked out cells were then grown for 2 to 3 days and seeded for the experiment. Cells were similarly electroporated with 1.5 nmol *JunB* guide RNA for activation, grown until confluency for 2 days and treated with 0 ng/ml or 400 ng/ml LPS without media removal. The *ETFDH* gene knockout or *JunB* gene activation were examined through western blot.

### Metabolomics

Metabolomics analysis was performed as previously described by Molenaars *et al.* 2021^28^. Dermal fibroblasts from both VLCADD patients, MADD patients and control individuals were seeded in triplicates in 35mm cell culture dishes with a density of 150,000 cells/dish in Ham’s F-10 nutrient mixture (Gibco) supplemented with L-glutamine (Bio-Whittaker), 10% foetal bovine serum (FBS) (Bio-Whittaker), 100 U/ml penicillin, 100μg/ml streptomycin (Life Technologies), and 250 ng/ml Fungizone (Life Technologies) in a humidified atmosphere with 5% CO2 at 37 °C. After 24 hours medium was changed to Ham’s F-10 nutrient mixture, 1% FBS (as described above) with 0 ng/mL or 400 ng/mL LPS (Serotype EH100 (RA) (TLRGRADE®) (Ready-to-Use), Alexis Biochemicals) and incubated for 24 hours (5% CO_2_ at 37°C). On ice, medium was removed and stored at -20°C before the following solvents were added to each well: 500 μL methanol, 425 μL Milli-Q, and 75 μL internal standard solution (in house solution, the full list available in Supplementary Materials). Cells were scraped off the dishes and both cells and solvents were pipetted into a 2 mL Eppendorf Safe-lock tube and stored at -80°C. Samples were prepared for analysis by adding 1 mL chloroform followed by 1.5 minutes of thorough mixing, and subsequent centrifugation at 20000 x g, 4°C for 10 minutes. Top phase was transferred to a 1.5 mL Eppendorf tube and dried using a vacuum concentrator at 60°C. Pellets were reconstituted in 100 μL 3:2 methanol: Milli-Q and samples were mixed thoroughly, before centrifugation at 20000 x g, 4°C for 10 minutes. 85 μL of each sample were transferred to a glass vial and stored at -20°C until mass spectrometry analysis. Metabolites were analysed using a Waters Acquity ultra-high performance liquid chromatography system coupled to a Bruker Impact II™ Ultra-High Resolution Qq-Time-Of-Flight mass spectrometer. Samples were kept at 12°C during analysis and 5 µL of each sample was injected. Chromatographic separation was achieved using a Merck Millipore SeQuant ZIC-cHILIC column (PEEK 100 x 2.1 mm, 3 µm particle size). Column temperature was held at 30°C. Mobile phase consisted of (A) 1:9 (v/v) acetonitrile: water and (B) 9:1 (v/v) acetonitrile: water, both containing 5 mmol/L ammonium acetate. Using a flow rate of 0.25 mL/min, the LC gradient consisted of: 100% B for 0-2 min, reach 54% B at 13.5 min, reach 0% B at 13.51 min, 0% B for 13.51-19 min, reach 100% B at 19.01 min, 100% B for 19.01-19.5 min. Equilibrate column by increasing flow rate to 0.4 mL/min at 100% B for 19.5-21 min. MS data were acquired using negative and positive ionization in full scan mode over the range of m/z 50-1200. Data were analysed using Bruker TASQ software version 2021.1.2 452. All reported metabolite intensities were normalized to total protein content in samples, determined using a Pierce^TM^ BCA Protein Assay Kit, as well as to internal standards with comparable retention times and response in the MS. Metabolite identification was based on a combination of accurate mass, (relative) retention times and fragmentation spectra, compared to the analysis of a library of standards^28,29^.

### Statistical analyses

For all experiments, biological cell culturing duplicates or triplicates have been included, indicated for each individual experiment in the figure legends. For some experiments, indicated in figure legends, also technical replicates have been included in the statistical analysis. Statistical significance between treated and non-treated individual cell lines are calculated using unpaired Student’s t-test. All statistical analyses have been performed using GraphPad Prism 9 (GraphPad Software, Inc). Not significant (not indicated): p≥0.05, *: p=0.01 to 0.05, **: p=0.001 to 0.01, ***: p<0.001, and ****: p<0.0001.

## RESULTS

### Deleterious *ETFDH* disease*-*variants of MADD patients cause impaired responses to LPS

We investigated the cytokine response to LPS using fibroblasts derived from two patients each carrying deleterious variants of *ETFDH,* ecoding the ETF-ubiquinone oxidoreductase (ETF-QO (Suppl. table 1)^30,31^. The patient variants are essentially null-variants as the ETFDH-encoded product ETF-QO is not expressed in the patient derived cells (Fig. 1A). In short, patient-derived fibroblasts (P1+P2) and fibroblasts derived from healthy controls (C1+C2) were stimulated with 400 ng/ml LPS. Cell supernatants were then collected to determine secretion of common pro-inflammatory cytokines IL-6 and IL-8 by ELISA. Here, fibroblast from healthy donors displayed significant LPS-induced release of these cytokines into the supernatant. By contrast, the LPS-induced cytokine release was markedly attenuated in fibroblasts derived from MADD patients (Fig. 1B-C and Suppl.Fig. 1A). This also seemed to be the case when measuring cytokine induction at the mRNA level by qPCR for *IL-6*, *IL-8*, *CCL2*, *IL-1β*, and *IL-1α,* (Fig. 1D-H). The response was not restricted to LPS as induction of CXCL10 was also attenuated in cells from MADD patients when stimulated with RNA in the form of poly I:C (Suppl. Fig. 1B). Because the two included MADD patients carry genetically distinct deleterious variants of *ETFDH* (Table 1), it is highly plausible that the shared inability to respond to LPS originates from their deficiencies in *ETFDH*. Nevertheless, we wanted to explore further, if *ETFDH* deficiency in itself is sufficient to suppress cytokine responses to LPS thus linking the known genotype directly to the phenotype. For this purpose, we silenced *ETFDH* expression using siRNAs in fibroblasts derived from healthy donors. Here, treatment with siRNA reduced the expression of ETF-QO but also reduced the induction of *IL-6, CCL2, CXCL10*, and *IL-1β* in response to stimulation with LPS as compared with cells treated with control siRNA (Figure 1I-O). These experiments support that the *ETFDH* deficiencies characterizing the MADD patients are sufficient to cause the insufficient response to LPS.

**Figure 1.**
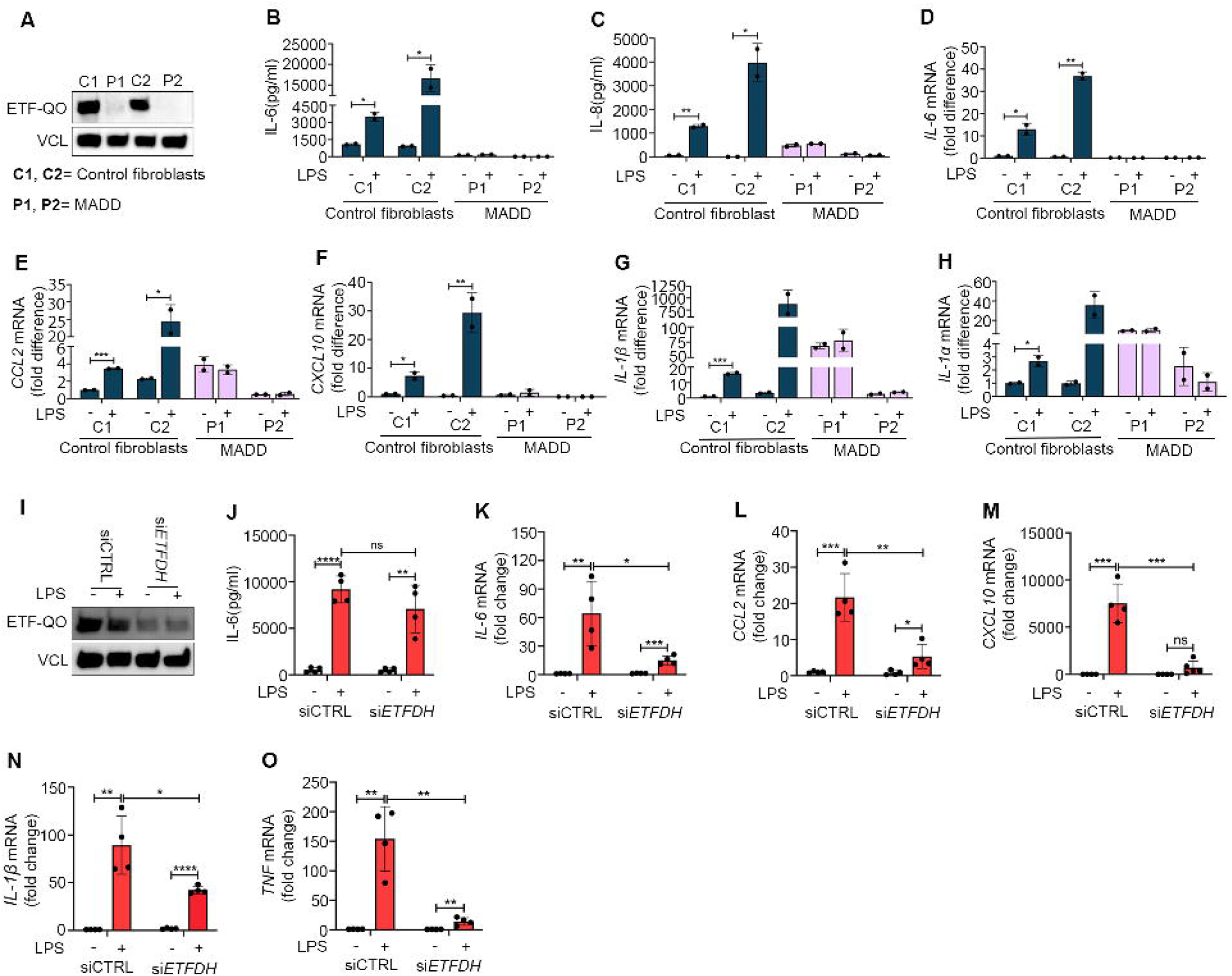
Inborn errors in *ETFDH* cause impaired response to LPS. Primary dermal fibroblasts from healthy controls (C1-2) and primary dermal fibroblasts derived from MADD patients (P1-2) were (A) blotted for basal level expression of ETF-QO. The cells were stimulated with LPS 400 ng/ml, and analyzed for (B-C) IL-6, IL-8 secretion at 24 h by ELISA or (D-H) cytokines mRNA (IL-6, CCL2, CXCL10, IL-1β, and IL-1α) expression by RT-QPCR at 6 h. Control dermal fibroblasts (C2) transfected with scrambled siRNA or *ETFDH* siRNA for 72 h were stimulated with LPS 400 ng/ml for (I) 24 h, blotted for ETF-QO, and analyzed for IL-6 secretion or (K-O) 6 h, and analyzed for cytokines mRNA (IL-6, CCL2, CXCL10, IL-1β, and TNF) expression by RT-QPCR. Blots in figures A and I are representative of three independent experiments. Data in figures B-D are representative of two independent experiments performed in biological duplicates. Data in figures E-H represent mean ± SEM of an experiment performed in biological duplicates. Data in figures J-O represent average mean ± SEM of two independent experiments performed in biological duplicates. *P < 0.05, **P < 0.01 ***P < 0.001, and ****P < 0.0001; compared to untreated control cells (unpaired t test). MADD: Multiple Acyl-CoA Dehydrogenase Deficiency; ETFDH: Electron Transfer Flavoprotein Dehydrogenase.

### MADD patient cells have reduced TLR4 expression and signaling

As the cytokine response of MADD cells with *ETFDH* deficiency to stimulation with LPS was impaired, we speculated that components of the TLR4 signaling pathway could be affected (Fig. 2A). Indeed, cells from MADD patients displayed markedly reduced *TLR4* expression levels when compared to cells from healthy human controls (Fig. 2B). Because of the reduced expression of TLR4 we investigated the proximal signaling event of Interleukin-1 Receptor-Associated-Kinase 1 (IRAK1) phosphorylation (pIRAK1). Here, pIRAK1 seemed to be reduced both at basal level as well as in response to LPS (Fig. 2C). By contrast, pIRAK1 levels were clearly increased in response to stimulation with IL-1β suggesting that IRAK1 is functional in these cells (Fig. 2D). Interestingly, the treatment of cells from healthy controls with siRNA targeting *ETFDH* also resulted in reduced expression of *TLR4* (Fig. 2E) and in reduced pIRAK1 levels (Fig. 2F).

**Figure 2.**
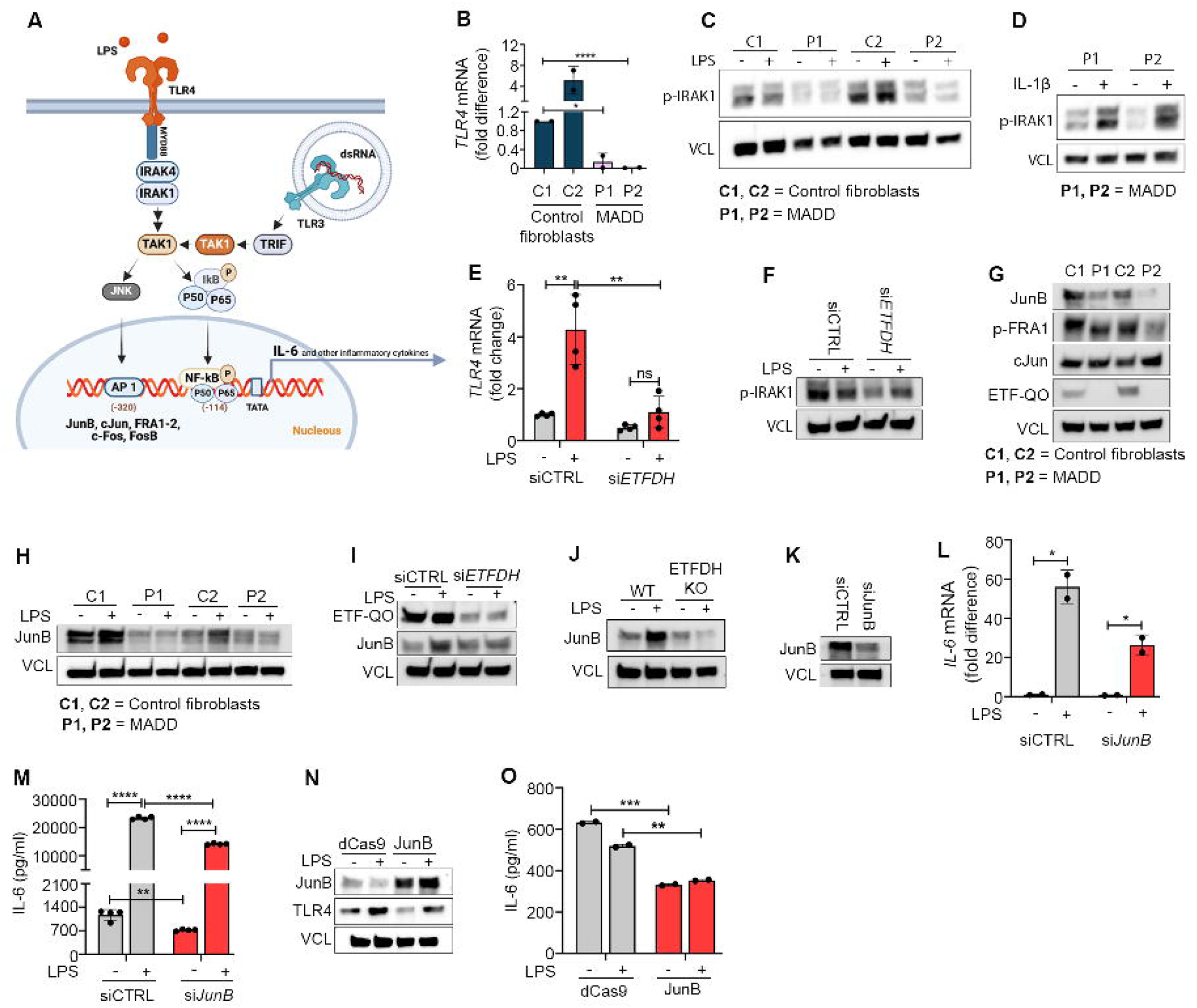
MADD patient cells have reduced TLR4 expression and signaling. (A) Selected proteins, signaling molecules and transcription factors involved in TLR4 and TLR3 signaling pathways. Primary dermal fibroblasts from healthy controls (C1-2) and primary dermal fibroblasts derived from MADD patients (P1-2) were (B) analyzed for basal TLR4 mRNA expression by RT-QPCR, (C-D) stimulated with LPS 400 ng/ml or recombinant human IL-1β 100 pg/ml for 15 min and blotted for phospho-IRAK1. The cells were (G) also blotted for basal expression of AP-1 transcription factors including JunB, phospho-FRA1, and cJun, or (H) stimulated with LPS 400 ng/ml for 24 h and blotted for JunB. Primary dermal fibroblasts from healthy controls (C2) transfected with scrambled siRNA or ETFDH siRNA for 72 h were stimulated with LPS 400 ng/ml and (E) analyzed for TLR4 mRNA TLR4 mRNA expression at 6 h, (F) blotted for phospho-IRAK1 at 2 h, and (I) blotted for ETF-QO and JunB at 24 h. (J) Control dermal fibroblasts (C2): both wild type (WT) and those electroporated with ETFDH sgRNA for ETF-QO knockout, were stimulated with LPS 400 ng/ml for 24 h and blotted for ETF-QO and JunB. (K-M) Primary dermal fibroblasts from healthy control (C2) transfected with scrambled siRNA or JunB siRNA for 72 h and stimulated with LPS 400 ng/ml. Cells were analyzed for IL-6 mRNA expression after 6 h and IL-6 secretion after 24 h. (N-O) MADD patients derived dermal fibroblasts (P1) were electroporated with dCas9 and JunB guide RNA for CRISPR activation. Collected cells were seeded and the confluent cells after 2-3 days were stimulated with LPS 400 ng/ml for 24 h, blotted for JunB, and analyzed for IL-6 secretion. Data in figures B, E, L represent mean ± SEM of two independent experiments performed in biological duplicates. Blots C-D and F-I are representatives of at least two independent experiments, while blots in figures J and N were performed once. Data in figure O, represent values of technical replicates from each case of figure N. *Represents significance compared to healthy controls or untreated control cells (*P < 0.05, **P < 0.01 ***P < 0.001, and ****P < 0.0001, unpaired t test).

We then decided to probe the expression of other TLR4-signaling components besides TLR4 itself and IRAK1. The hetero-dimeric transcription factor AP1 is crucial for IL-6 induction and TLR4-induced transcriptional activation. AP1 functions as a dimer in different combinations with members of the Jun family (c-Jun, JunB, and JunD) and the Fos family (c-Fos, FosB, Fra1, and Fra2)^32^. Here, we could observe a notably reduced expression of JunB as well as slightly reduced expression levels of cFOS and CREB in cells from MADD patients whereas c-Jun was found to be equally expressed between patient and control cells (Fig. 2G and Supp. Fig. 2). We further investigated the expression of JunB in controls as well as in MADD cells 24 h after LPS stimulation. Here, LPS stimulation led to increased expression of JunB in cells from controls (C1, C2, C3, and C4), whereas the expression was either below detection levels or failed to increase in response to LPS in MADD cells (P1, P2) (Fig. 2H). Furthermore, we found that inhibition of *ETFDH* by siRNA or by CRISPR/Cas9 mediated knock-out was sufficient to also reduce JunB expression levels hereby connecting reduced JunB expression and responsiveness to LPS to the genotype of MADD patients (Fig. 2I-J and Supp. Fig. 3). To test if suppression of JunB was sufficient to suppress IL-6 expression, we performed siRNA mediated JunB silencing in control cells. JunB silencing significantly decreased LPS induced IL-6 mRNA and IL-6 secretion. Importantly, JunB silencing also decreased basal level of IL-6 expression (Fig. 2K-L). This highlighted the crucial role of JunB in IL-6 response and infers that its reduced expression might be one of the responsible factors for the impaired IL-6 secretion in patient cells. This inspired us to activate JunB in patient cells to test if this would be sufficient to restore the IL-6 response. We performed CRISPR activation for JunB on MADD patient cells using specific guide RNA (gRNA) and nuclease-dead Cas9 (dCas9). We observed a significant increase in JunB expression upon CRISPR activation. However, neither TLR4 expression nor IL-6 induction was rescued by the forced expression of JunB, suggesting that although JunB is a contributing factor, additional factors are significant for the impaired LPS-response in MADD (Fig. 2M-O).

### Deleterious *ACADVL* disease*-*variants also cause impaired responses to LPS

To investigate if the observed inability to respond appropriately to stimulation with LPS was specific to MADD patients or a common trait in lcFAODs, we investigated fibroblasts from patients with VLCADD. This line of experiments included cells from patients with severe VLCADD (P5-P8), and with mild VLCADD (P3 and P4) expressing disease-causing variants of *ACADVL* (Suppl. table 1)^33–35^. Interestingly, we observed that similarly to MADD patient cells, release of IL-6 and IL-8 to the cell supernatant in response to stimulation with LPS was also attenuated in cells from VLCADD patients (Fig. 3A-B). This was also observed at the mRNA level where LPS did not induce significant expression of *IL-6.* In contrast to MADD patient cells, some of the VLCADD patients displayed some induction of *CCL2* and IL-1α at the mRNA level (Fig. 3C-F). As with the MADD deficiencies, we then tried to establish if the deficiencies in *ACADVL* could be linked directly to the insufficient response to LPS. We therefore treated fibroblasts derived from healthy human controls with siRNA specific for *ACADVL* or with control siRNA (Fig. 3G). Again, siRNA-induced deficiency of *ACADVL* led to decreased LPS-induced release of IL-6 and IL-8 to the supernatant (Fig. 3H-I). This effect was also observed at the mRNA level with reduced induction of *IL-6, CCL2*, *IL-1β*, *IL-1α, CXCL10, and TNFα* (Fig. 3J-O). Similar to the *ETFDH,* siRNA targeting *ACADVL* resulted the reduced TLR4 mRNA expression (Fig. 3P). Importantly, as was also observed with cells from MADD patient, TLR4 expression levels were reduced when comparing VLCADD patient cells with cells from healthy controls (Fig. 3Q-R). Also, LPS stimulation induced expression of JunB in the control fibroblasts (C3, C4), while in the VLCADD cells (P5, P8), the expression was either not observed or it was not induced (Fig 3S). To further test the linkage between VLCADD and insufficient responses to LPS, we stimulated cells from two patients presenting mild forms of VLCADD. Interestingly, the LPS-response was less attenuated in these patients with only some repression of IL-6 responses and clear inductions of IL-8 (Suppl. Fig 4A-B). In addition, TLR4 expression levels were markedly higher in cells from mild VLCADD patients when compared to cells from patients presenting with severe VLCADD (Suppl. Fig. 4C-D).

**Figure 3.**
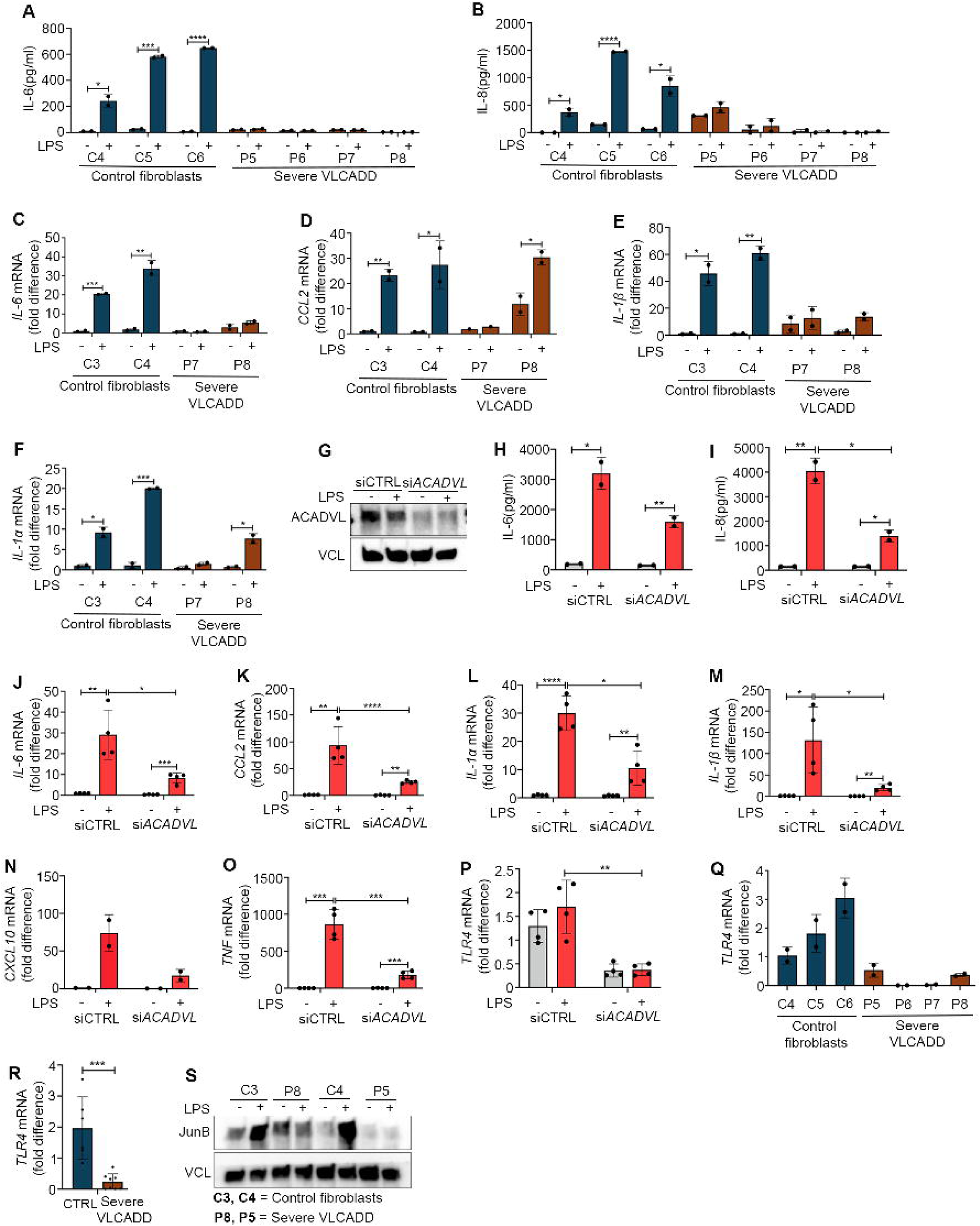
Reduced LPS responsiveness in VLCADD patient cells. Primary dermal fibroblasts from healthy controls (C4-6) and primary dermal fibroblasts derived from severe VLCADD patients (P5-8) stimulated with LPS 400 ng/ml were analyzed for (A-B) IL-6, IL-8 secretion at 24 h and (C-F) cytokines mRNA (IL-6, CCL2, IL-1β, and IL-1α) expression by RT-QPCR at 6 h. (G-P) Primary dermal fibroblasts from healthy controls (C4) transfected with scrambled siRNA or *ACADVL* siRNA for 72 h were stimulated with LPS 400 ng/ml and blotted for ACADVL, analyzed for cytokines secretion (IL-6, IL-8) at 24 h, cytokines mRNA (IL-6, CCL2, IL-1α, IL-1β, CXCL10, TNF), and TLR4 mRNA expression by RT-QPCR at 6 h. (Q-R) Primary dermal fibroblasts from healthy controls (C4-6) and primary dermal fibroblasts derived from severe VLCADD patients (P5-8) were analyzed for basal level TLR4 mRNA expression. (S) The cells were stimulated with LPS 400 ng/ml for 24 h and blotted for JunB. Data in figures A-F represent mean ± SEM of one experiment performed in biological duplicates. Data in figures H-P represent average mean ± SEM of two independent experiments performed in biological duplicates. Blots in figures G and S were performed twice in independent experiments. *Represents significance compared to healthy controls or untreated control cells (*P < 0.05, **P < 0.01, ***P < 0.001, and ****P < 0.0001, unpaired t test).

Finally, in contrast to the MADD cells, the cytokine response to RNA in the form of poly-I:C did not seem to be affected in cells from either mild or severe forms of VLCADD (Suppl. Fig. 5). These data demonstrate a clear link between genes that are important for beta-oxidation and LPD-induced inflammatory responses.

### Pharmacological blockage of fatty acid oxidation does not impair the IL-6 response

Cells from MADD and VLCADD patients have severely compromised lcFAO as demonstrated by reduced lcFAO flux (Figure 4A-C). Inhibition of *ETFDH* by siRNA recapitulated this reduction in lcFAO (Figure 4D). We therefore speculated that inhibition of lcFAO using etomoxir (ETO), which blocks FAO by inhibiting influx of fatty acids from the cytosol into the mitochondria by direct inhibition of CPT1, would affect responsiveness to LPS. Although ETO clearly abrogated lcFAO flux, it did not affect the ability of cells from healthy controls to respond to LPS with secretion of IL-6 (Figure 4E-F), nor did it affect the already reduced IL-6 expression in patients’ cells. These results imply that the impaired response to LPS in cells with genetic deficiencies in *ETFDH* or *ACADVL* might not be due to the blockage of lcFAO per se.

**Figure 4.**
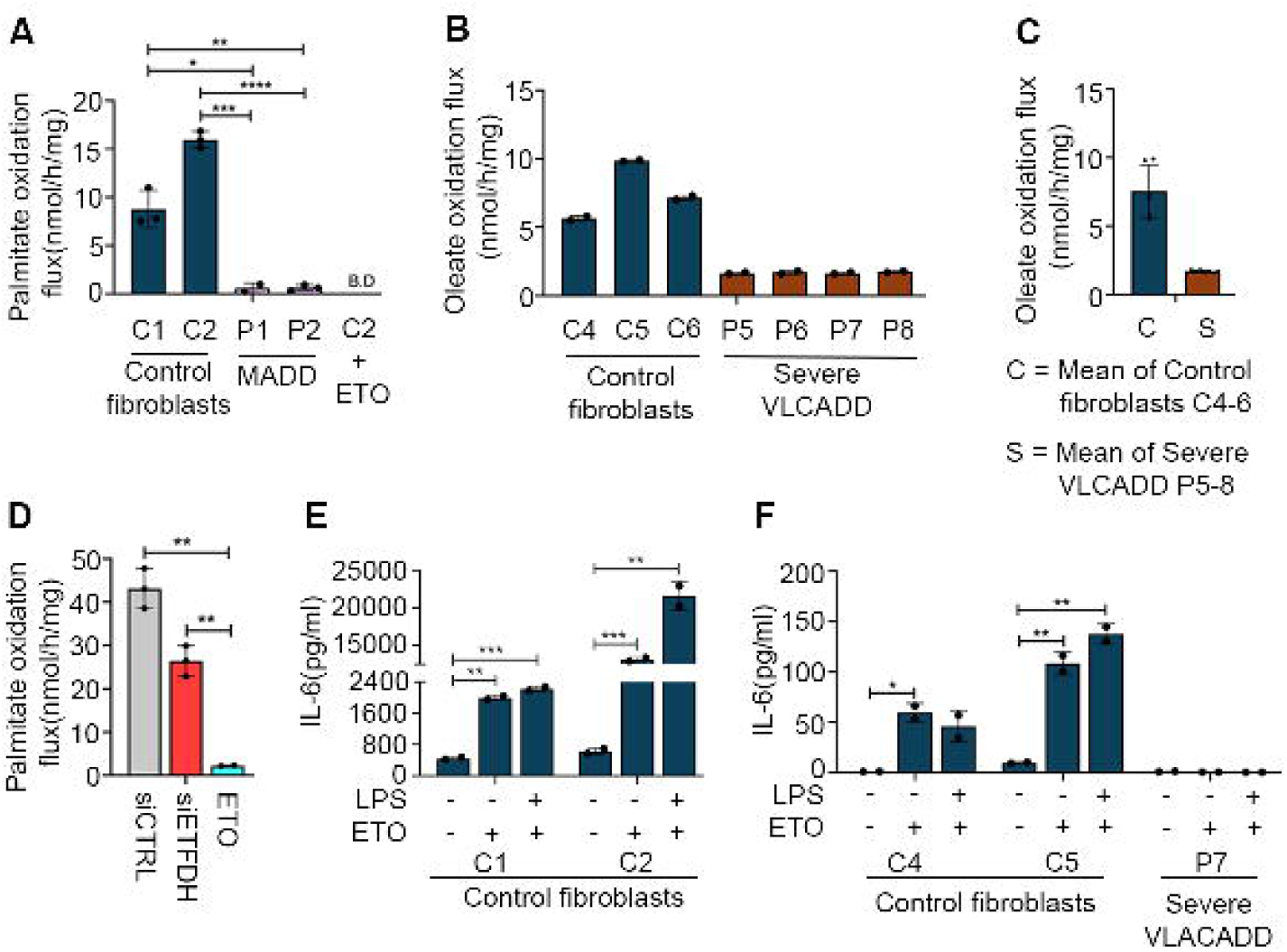
Pharmacological blockage of fatty acid oxidation does not impair cytokine responses to LPS. (A) Palmitate oxidation flux of primary dermal fibroblasts from healthy controls (C1-2) and dermal fibroblasts derived from two MADD patients (P1-2) at 37°C. Control fibroblast cells (C2) were preincubated with Etomoxir (ETO) overnight before measurement as a negative control. (B) Oleate oxidation flux of control dermal fibroblasts (C4-6) and dermal fibroblast cells from severe VLCADD (P5-8) measured at 37°C. (C) Group means calculated from data in B, showing overall FAO flux differences in control dermal fibroblasts (C4-6) and severe VLCADD (P5-8). D) Primary dermal fibroblasts from healthy controls C2 were transfected with scrambled siRNA or ETFDH siRNA for 72 h, or preincubated with ETO overnight before palmitate oxidation flux measurement. (E and F) Primary dermal fibroblasts from healthy controls (C1-2, C4-5) and severe VLCADD (P7), overnight incubated with ETO were further stimulated with LPS 400 ng/ml for further 24 h before measuring IL-6 by ELISA. Data are presented as mean ± SEM; number of cell culture replicates: 2 to 3. Each cell culture had 2 to 3 (FAO flux) or 2 (ELISA) technical replicates. *P < 0.05, **P < 0.01, ***P < 0.001; compared to healthy controls or untreated control cells (unpaired t test).

### Inhibition of mTOR with rapamycin partially restores basal IL-6 secretion but not responses to LPS

In order to explore potential drivers of the impaired cytokine response in severe VLCADD and MADD, we performed intracellular metabolomics analyses of dermal fibroblasts from controls, severe VLCADD, mild VLCADD, and MADD, incubated both with and without the addition of LPS, which were subsequently analyzed by UPLC-mass spectrometry. We detected 112 unique metabolites, encompassing classes such as amino acids, glycolysis and TCA cycle intermediates, and nucleotides. Metabolomics analysis detected significant increases in essential and non-essential amino acids, both at basal level and after LPS stimulation in dermal fibroblasts from severely affected VLCADD subjects, compared to control dermal fibroblasts (Suppl. Fig. 6A-B). Untreated severe MADD dermal fibroblasts also show an increase in some amino acids compared to control dermal fibroblasts. Unlike VLCADD, MADD cells show a clear decrease in the abundance of several metabolites, including various amino acids, after LPS stimulation (Suppl. Fig. 6B).

Using the metabolic aberrations in patient cells as a starting point, we searched for a number of ways that could be helpful to rescue IL-6 levels in patient cells. Severe VLCADD cells showed significant accumulation of several essential and non-essential amino acids, compared to controls, both with or without LPS stimulation. Multiple amino acids are known to influence mTOR signaling by enhancing RAG proteins activation that are essential in recruiting mTOR^36^. It has been shown that mTOR pathway activation limits NF-kB activity by enhancing IkBα and thereby suppressing proinflammatory markers in fibroblasts^37^. Therefore, we wanted to test if mTOR inhibition could be sufficient to fully or partially restore IL-6 level in patient cells only, where mTOR inhibition is expected. MADD cells (P1) and VLCADD cells (P8) were preincubated with the well-known inhibitor of mTOR, rapamycin (100 nM) for 24 h, followed by LPS stimulation for a further 24 h without media removal. Rapamycin preincubation decreased phosphorylated mTOR expression and we observed significant increase in basal level IL-6 secretion in patient cells. However, rapamycin did not restore responsiveness to stimulation with LPS (Suppl. Fig 6C-E). To further test the involvement of the mTOR pathway we performed *MTOR* silencing by siRNA in MADD cells (P1, P2). Blotting for phospho-mTOR showed significantly decreased expressions in si*MTOR* conditions when compared to scrambled siRNA, demonstrating that silencing was effective. Cells transfected with si*MTOR* showed a significantly larger basal level increase in IL-6 secretion than the respective controls with scrambled siRNA transfection. However, as with rapamycin, responsiveness to LPS was not restored (Suppl. Fig 7F-H). Our results imply that mTOR cold be involved in repressing IL-6 expression at the basal level but that inhibition of mTOR is not sufficient to restore LPS sensitivity.

## DISCUSSION

With this work, we provide genetic evidence from human patient samples, that enzymes involved in lcFAO are important for responsiveness to LPS. We observed this phenomenon across six genetically distinct individuals – two MADD patients with distinct *ETFDH* deficiencies and four VLCADD patients with distinct *ACADVL* deficiencies. Our results therefore strongly suggest that a connection between lcFAO enzymes and inflammatory responses exists. This was corroborated by the observation that genetic loss-of-function of *ETFDH* and *ACADVL* through siRNA in fibroblasts from healthy subjects generated a similar phenotype.

Dysregulated immune responses in lcFAOD were recently reported in both animal models as well as patients^17,21,22,38^, however, the underlying molecular mechanisms remain undiscovered. Further, several studies investigating insulin resistance and obesity have reported that long-chain saturated fatty acids such as palmitate and long-chain acylcarnitines can serve as inflammatory triggers. However, the target through which long-chain acylcarnitines activate an inflammatory response is not yet known^39^. In this study we learned that blocking lcFAO by etomoxir did in itself not affect responses to LPS indicating that an effect of lcFAOD besides inability to metabolize fatty acids is likely part of the underlying mechanism. These data therefore also suggest that FAO is not a driver of IL-6 induction. Dysfunctional response to LPS in leukocytes has recently also been observed in respiratory chain disorders. Karan *et al.* proposes that cytokine responses are blunted in patients with mtDNA deletions encoding respiratory chain proteins^40^. Patients with respiratory chain disorders often have overlapping clinical and biochemical phenotypes with MADD patients^41–43^.

TLR4 is an important member of the toll-like receptor family and is broadly expressed in human cells and tissue, i.e., liver, skin, heart, muscle and adipocytes^44–46^. Our studies demonstrate that cells derived from patients with lcFAOD, with permanent blockage of lcFAO have constantly decreased TLR4 levels. Further, we link decreased expression of important transcription factors that can interact with the IL-6 promoter for mRNA synthesis to the impaired IL-6 response. This could possibly explain the lack of basal IL-6 response in patients’ cells and also provides an opportunity to restore them.

Together, this study links inborn errors in long-chain fatty acid oxidation to insufficient responses to LPS.

## SUPPLEMENTARY FIGURE LEGENDS

**Supplementary figure 1. Cytokine responses in MADD.**

Primary dermal fibroblasts from healthy controls (C1-2) and primary dermal fibroblasts derived from MADD (P1-2) were stimulated with LPS 400 ng/ml for 24 h or preincubated overnight with poly IC 2 µg/ml, and analyzed for CXCL10 and IL-6 secretion by ELISA respectively. Data presented as mean ± SEM, and representative of two independent experiments performed in duplicates. *Represents significance compared to untreated control cells (***P < 0.001, and ****P < 0.0001, unpaired t test).

**Supplementary figure 2. IL-6 promoter linked transcription factors upon LPS stimulation.**

Primary dermal fibroblasts from healthy controls (C1-2) and MADD fibroblasts (P1) stimulated with LPS 400 ng/ml were blotted for IL-6 promoter linked transcription factors expressions. Blots were performed twice in the independent experiments.

**Supplementary figure 3. ETF-QO knockout in control fibroblasts.**

Primary dermal fibroblasts from healthy controls (C2) were electroporated with ETFDH sgRNA for ETF-QO knockout and the cells were blotted for ETF-QO, along with wild type (WT) cells. Knockout was performed as single and double knockout in a single experiment.

**Supplementary figure 4. Mild VLCADD patient cells show LPS responsiveness.**

Primary dermal fibroblasts derived from mild VLCADD patients (P3-4) and severe VLCADD patients (P5-8) stimulated with LPS 400 ng/ml were analyzed for (A-B) IL-6, IL-8 secretion at 24 h and (C) TLR4 mRNA expression by RT-QPCR at 6 h. (D) Group means calculated from data in C, showing representative TLR4 mRNA expression. Data presented as mean ± SEM from an experiment performed in a biological duplicate, which is also repeated in another independent experiment. *Represents significance compared to untreated control cells (**P < 0.01, ***P < 0.001, and ****P < 0.0001, unpaired t test).

**Supplementary figure 5. Mild and severe VLCADD patient cells show proper responses to polyIC.**

Primary dermal fibroblasts from healthy controls (C5-6) and primary dermal fibroblasts derived from mild VLCADD (P3-4), and severe VLCADD (P5-8) patients’ cells were preincubated overnight with poly IC 2 µg/ml and analyzed for IL-6 secretion by ELISA. Data presented as mean ± SEM from an experiment performed in biological triplicates. ****P < 0.0001; compared to untreated control cells (unpaired t test).

**Supplementary figure. 6. Reduced IL-6 responses in MADD and VLCADD involves mTOR.**

(A-B) Volcano plot of metabolomics performed using MS/MS. (A) Control dermal fibroblasts (N=4, C3-6) vs. severe VLCADD (N=4, P5-8). (B) Primary dermal fibroblasts from healthy controls (N=2, C1-2) vs. severe MADD (N=2, P1-2). (C – H) MADD (P1-2) and VLCADD (P8) patient derived dermal fibroblasts were preincubated with 100 nM rapamycin for 24 h, or transfected with scrambled siRNA and siMTOR for 72 h, and stimulated with LPS 400 ng/ml for 24 h. Cells were then analyzed for IL-6 secretion and blotted for phospho mTOR. Data are presented as mean ± SEM of an experiment with 3 to 4 ELISA technical replicates. Experiment has been performed twice independently. *Represents significance compared to untreated control cells (*P < 0.05, **P < 0.01, ***P < 0.001, and ****P < 0.0001; unpaired t test).

## SUPPLEMENTARY MATERIALS

**Internal standard solution content for metabolomics investigations:**

Internal standard solution containing: adenosine-15N5-monophosphate (5 nmol), adenosine-15N5-triphosphate (5 nmol), D4-alanine (0.5 nmol), D7-arginine (0.5 nmol), D3-aspartic acid (0.5 nmol), D3-carnitine (0.5 nmol), D4-citric acid (0.5 nmol), 13C1-citrulline (0.5 nmol), 13C6-fructose-1,6-diphosphate (1 nmol), guanosine-15N5-monophosphate (5 nmol), guanosine-15N5-triphosphate (5 nmol), 13C6-glucose (10 nmol), 13C6-glucose-6-phosphate (1 nmol), D3-glutamic acid (0.5 nmol), D5-glutamine (0.5 nmol), D5-glutathione (1 nmol), 13C6-isoleucine (0.5 nmol), D3-lactic acid (1 nmol), D3-leucine (0.5 nmol), D4-lysine (0.5 nmol), D3-methionine (0.5 nmol), D6-ornithine (0.5 nmol), D5-phenylalanine (0.5 nmol), D7-proline (0.5 nmol), 13C3-pyruvate (0.5 nmol), D3-serine (0.5 nmol), D6-succinic acid (0.5 nmol), D5-tryptophan (0.5 nmol), D4-tyrosine (0.5 nmol), D8-valine (0.5 nmol).

**Supplementary table 1.**
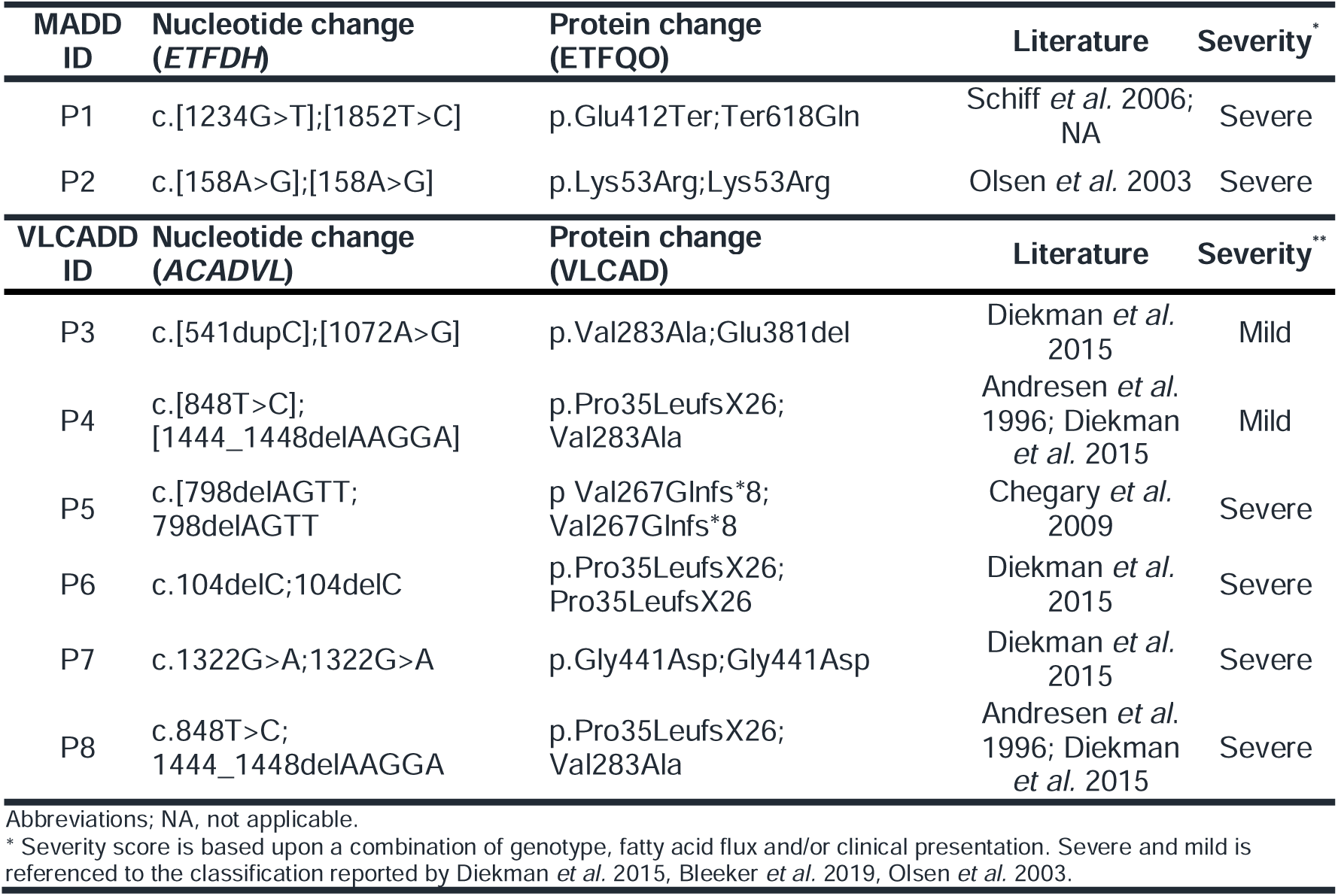
Patient cell lines genotypes.

**Supplementary table 2.**
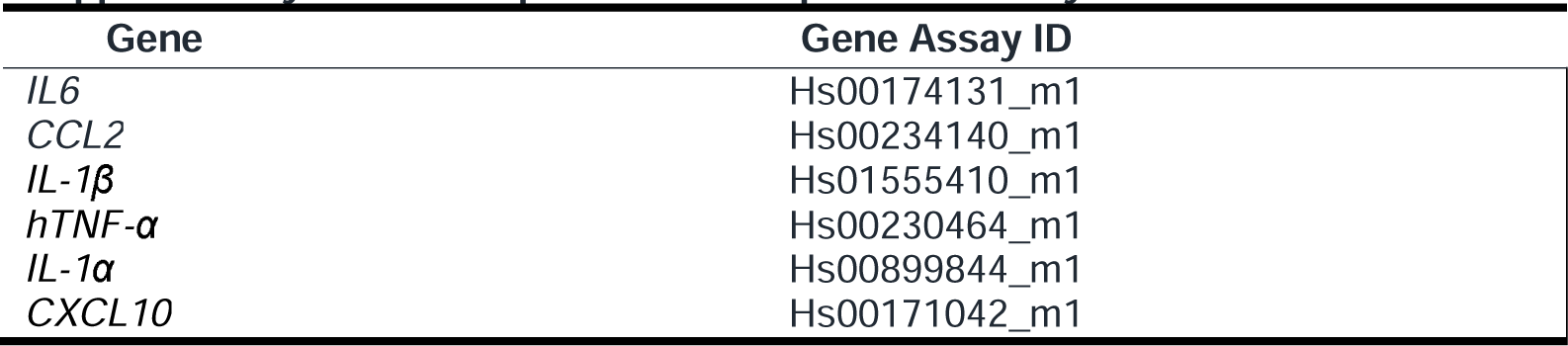
TaqMan® Gene Expression Assays.

**Supplementary table 3.**
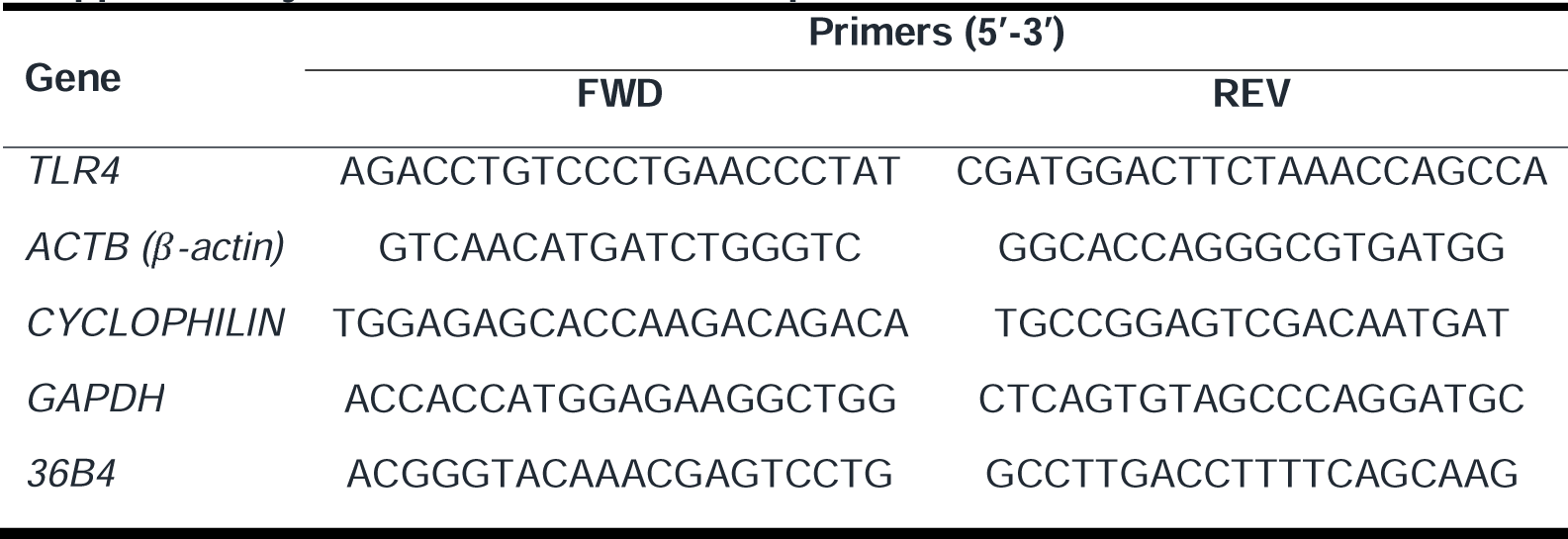
Quantitative PCR-primer sets.

