## Supplementary figures and images for "Human inborn errors of long-chain fatty acid oxidation show impaired inflammatory responses to TLR4-ligand LPS"

### Supplementary Figure 1

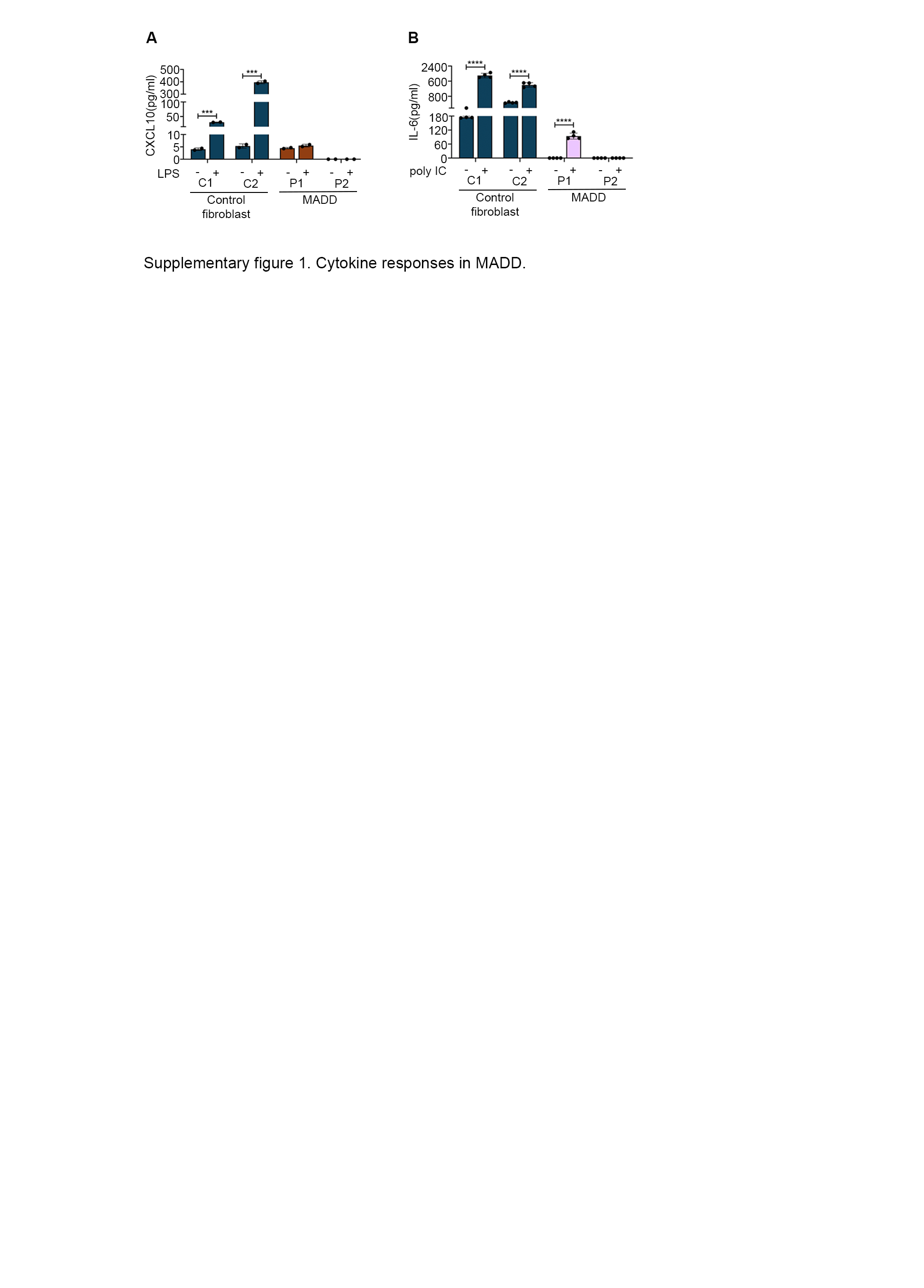

### Supplementary Figure 2 and 3

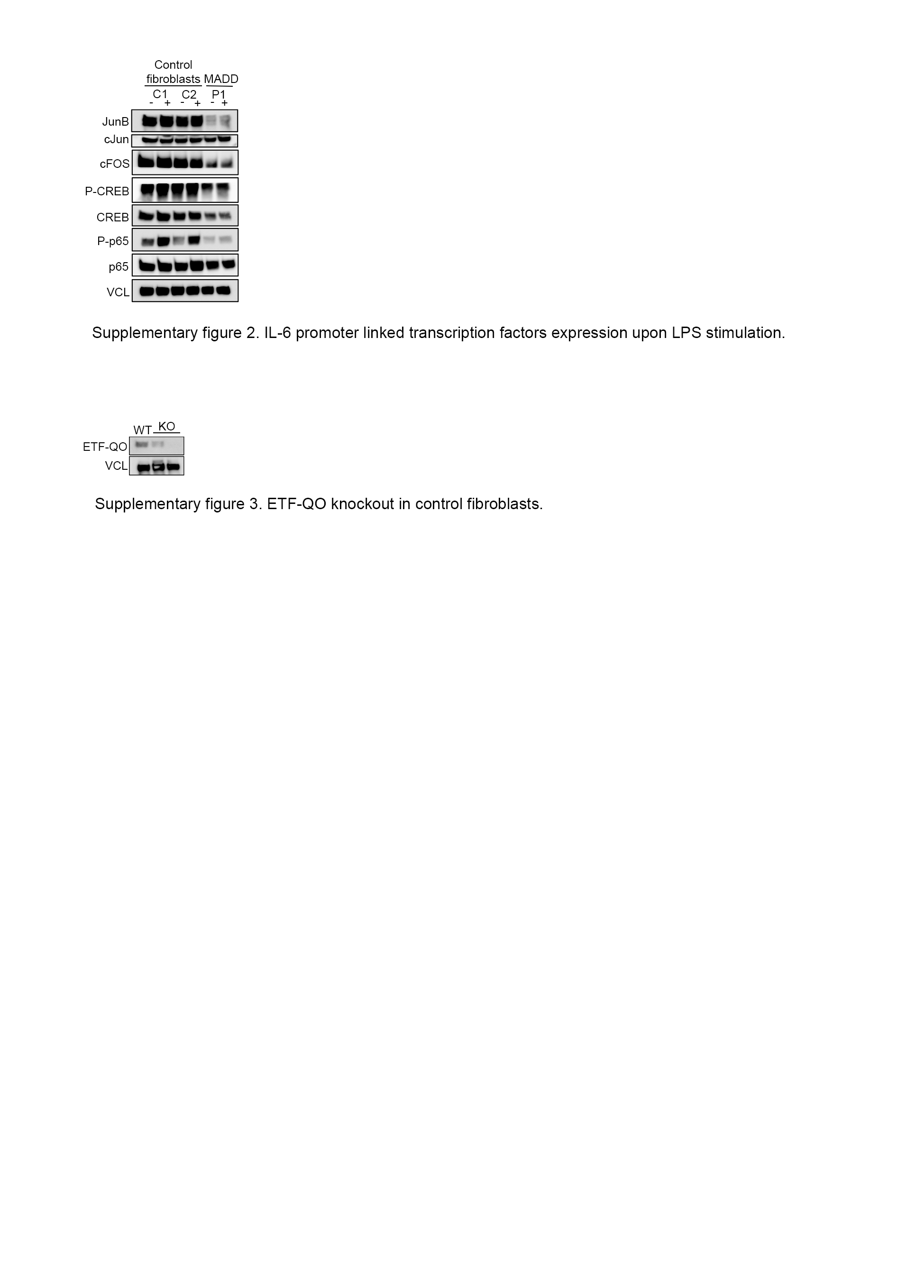

### Supplementary Figure 4 and 5

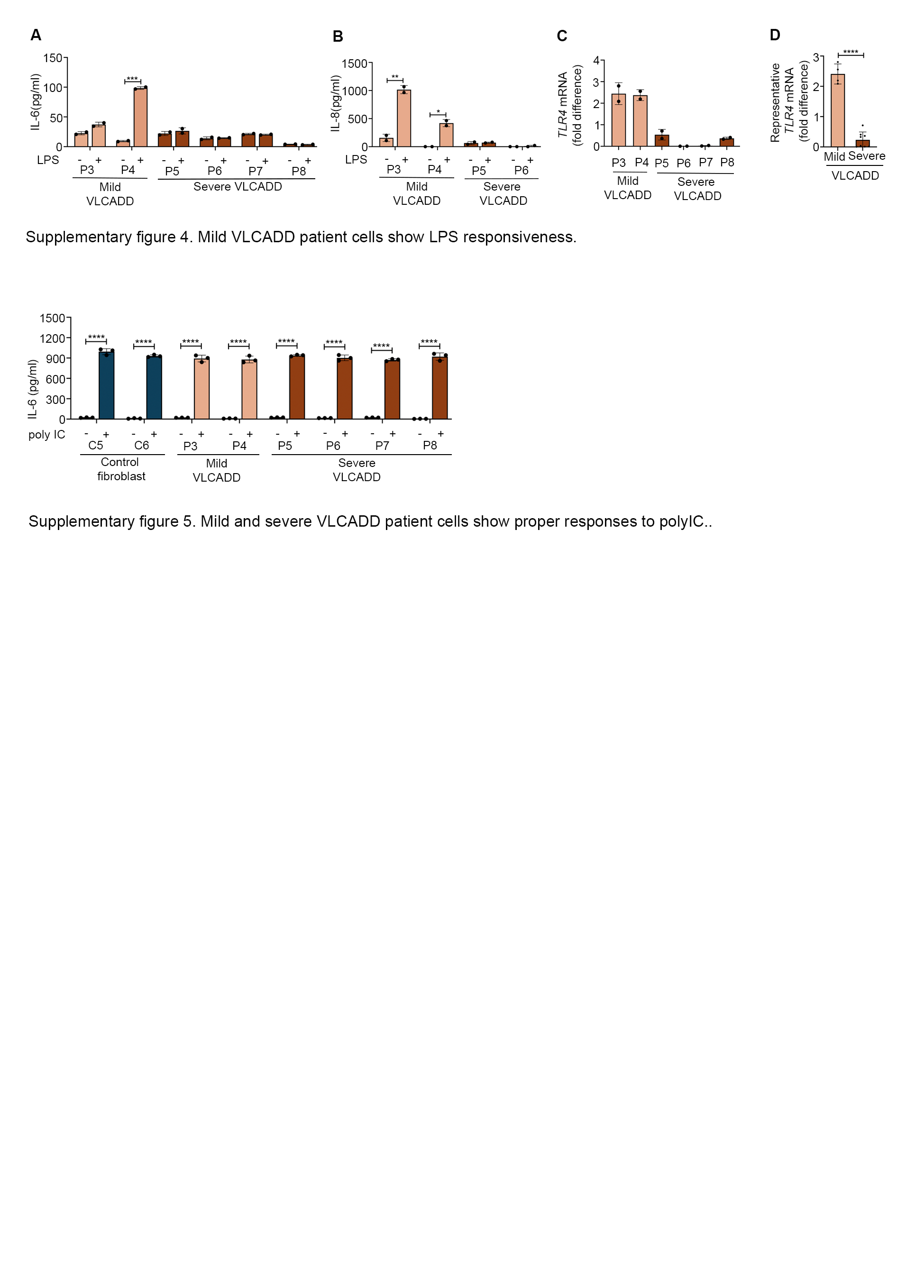

### Supplementary Figure 6

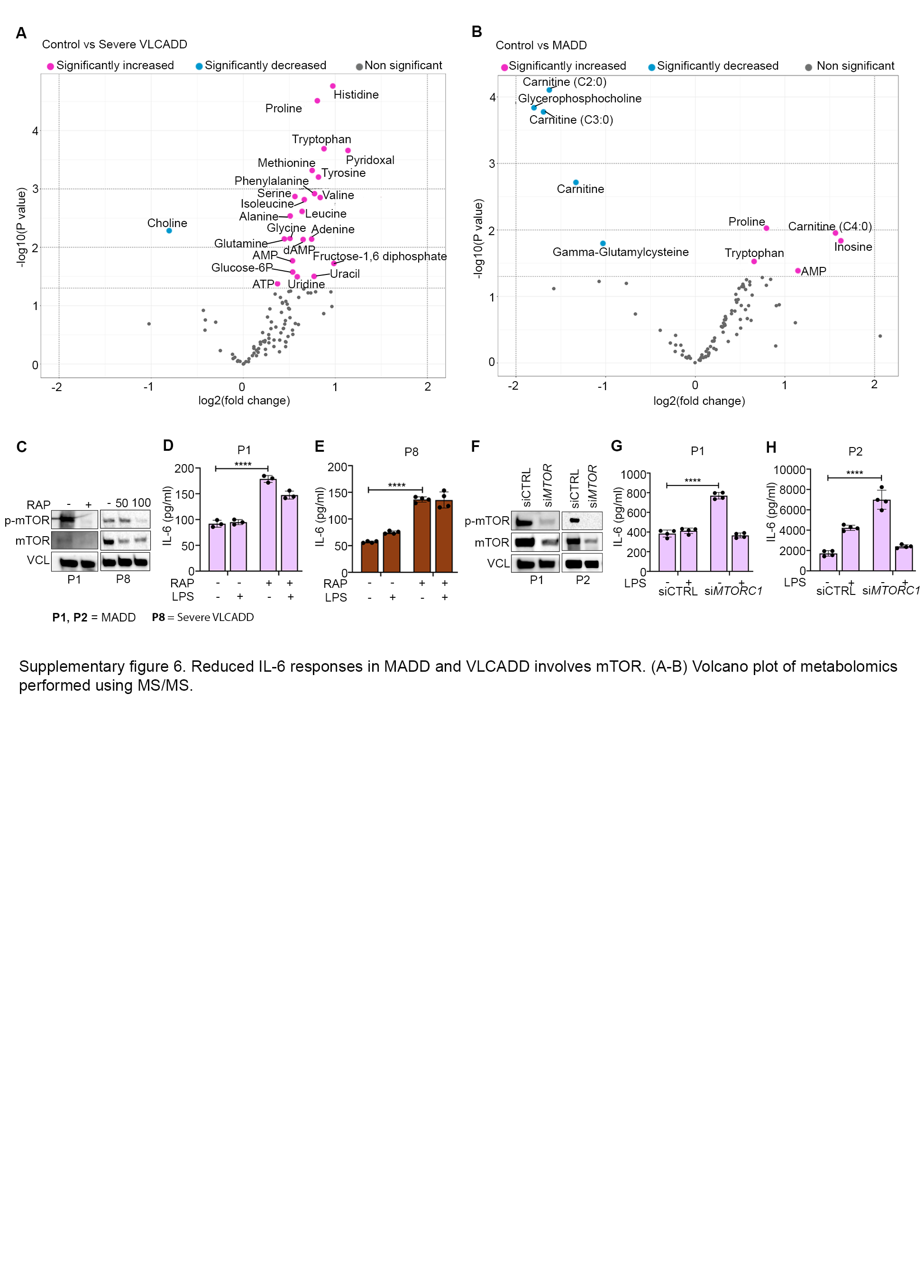
